# Persistent ventricle partitioning in the adult zebrafish heart

**DOI:** 10.1101/2021.03.16.435658

**Authors:** Catherine Pfefferli, Hannah R. Moran, Anastasia Felker, Christian Mosimann, Anna Jaźwińska

**Author notes:** equal contribution (Catherine Pfefferli, Hannah R. Moran).

## Abstract

The vertebrate heart integrates cells from the early-differentiating first heart field (FHF) and the later-differentiating second heart field (SHF) emerging from the lateral plate mesoderm. In mammals, this process forms the basis for the development of the left and right ventricle chambers and subsequent chamber septation. The single ventricle-forming zebrafish heart also integrates FHF and SHF lineages during embryogenesis, yet the contributions of these two myocardial lineages to the adult zebrafish heart remain incompletely understood. Here, we characterize the myocardial labeling of FHF descendants in both the developing and adult zebrafish ventricle. Expanding previous findings, late gastrulation-stage labeling using *drl*-driven CreERT2 recombinase with a myocardium-specific, *myl7*-controlled *loxP* reporter results in predominant labeling of FHF-derived outer curvature and the right side of the embryonic ventricle. Raised to adulthood, such lineage-labeled hearts retain broad areas of FHF cardiomyocytes in a region of the ventricle that is positioned at the opposite side to the atrium and encompasses the apex. Our data add to the increasing evidence for a persisting cell-based compartmentalization of the adult zebrafish ventricle even in the absence of any physical boundary.

## 1. Introduction

The heart underwent various specializations and adaptions supporting the wide variety of vertebrate body plans. Studies on cardiac development, physiology, and regeneration across model organisms have nonetheless revealed genetic and developmental features that are common to all vertebrate hearts (Lambers and Kume, 2016; Swedlund and Lescroart, 2019; Yao et al., 2021). Driving systemic blood circulation, the zebrafish heart’s ventricle is devoid of any larger, physical compartmentalization. Midline migration of bilateral heart progenitor fields in the lateral plate mesoderm (LPM) results in the formation of a cardiac disc, cone, and continuously extruding heart tube between 18-24 hours post-fertilization (hpf) (Bakkers, 2011; Felker et al., 2018; Kemmler et al., 2021).

The developing zebrafish heart incorporates both early-differentiating myocardium at its center and late-differentiating myocardium at the arterial and venous poles that have been linked to distinct progenitor pools within the anterior LPM (ALPM), deemed the first versus the second heart fields (FHF and SHF, respectively) (de Pater et al., 2009; Hami et al., 2011; Lazic and Scott, 2011; Zhou et al., 2011). While the FHF-assigned progenitors form the first differentiated myocardium of the initial linear heart tube, the SHF descendants continuously extend the elongating heart tube, yet differentiate at a later stage. This distinct differentiation timing results in FHF myocardium that is functional by 24 hpf and SHF that continues to differentiate until approximately 2-3 days postfertilization (dpf) (de Pater et al., 2009; Hami et al., 2011; Lazic and Scott, 2011; Mosimann et al., 2015; Zhou et al., 2011). The sub-division into distinct myocardial progenitor pools is of ancient evolutionary origin and a feature of heart across chordates, yet its purpose remains unclear (Kelly, 2012; Kelly et al., 2014; Prummel et al., 2020; Tolkin and Christiaen, 2012).

In zebrafish, even in the absence of physical ventricle compartmentalization, perturbation of the relative contributions of FHF and SHF causes defects in heart looping, outflow tract (OFT) patterning, and electrophysiological coupling, emphasizing the existence of cell-level compartmentalization within the developing atrium and ventricle (de Pater et al., 2009; Guner-Ataman et al., 2013; Hami et al., 2011; Mosimann et al., 2015; Rydeen et al., 2016; Zeng and Yelon, 2014; Zhou et al., 2011). In mammals and birds, the ventricle separates into distinct left and right chambers through septation that follows increasing compartmentalization at the gene expression level between the FHF-derived left and the SHF-derived right ventricle (Buckingham et al., 2005; Kelly et al., 2014; Morgenthau and Frishman, 2017; Mori and Bruneau, 2004; Vincent and Buckingham, 2010). A gradient of Tbx5 has been linked to refining the septation emerging between the FHF and SHF domains of the developing ventricle, a key feature towards separating systemic versus pulmonary circulation (Hanemaaijer et al., 2019; Koshiba-Takeuchi et al., 2009). While zebrafish form a single ventricle for systemic circulation without any appreciable physical septation, how the distribution of FHF versus SHF myocardium resolves in the adult teleost ventricle remains uncertain.

Beyond the contribution of both FHF and SHF myocardium to the heart, an increasing number of findings have revealed additional cellular or gene-level compartmentalization or lateralization in the zebrafish heart at embryonic and adult stages. *tbx5a* expression and function contributes to asymmetrical convergence of the bilateral heart field, with preferential migration of myocardial progenitors from the righthand ALPM (Mao et al., 2015). Manifesting in mid to late somitogenesis, the expression of *meis2b* divides the zebrafish atrium into two domains along the anterior-posterior axis of the heart field-forming ALPM, with *meis2b*-expressing posterior heart field progenitors preferentially contributing to the left side of the atrium in later development and in the adult heart (Guerra et al., 2018). What other such lateralization occurs in the zebrafish heart, in particular the ventricle, and how they contribute to cardiac function remain to be discovered.

We previously identified distinct gene-regulatory elements in the zebrafish-specific *draculin* (*drl*) locus that enabled the generation of transgenic reporters labeling distinct aspects of LPM patterning (Mosimann et al., 2015). Expression of *drl* during gastrulation is broadly confined to the emerging LPM-primed mesendoderm as directed by an intronic enhancer that responds to LPM-determining cues in zebrafish and in a variety of chordate models (Mosimann et al., 2015; Prummel et al., 2019). Together with additional, later-acting regulatory elements (Mosimann et al., 2015; Prummel et al., 2019), the fluorescent transgenic *drl* reporter activity during somitogenesis increasingly refines to cardiovascular lineages in zebrafish, and in the heart predominantly to the FHF-derived endocardium and myocardium of the hearts (Felker et al., 2018; Mosimann et al., 2015; Sánchez-Iranzo et al., 2018). *drl*-based zebrafish transgenics provide genetic means to further elucidate FHF lineages and patterning dynamics across developmental stages.

The binary Cre/*lox* system is a widespread tool for genetic lineage labeling. Deployed in a variety of model organisms including in zebrafish, Cre recombinase enables cassette excision or inversion of *loxP*-flanked transgene cassettes, resulting in permanently changing transgenic effector or reporter expression in cells with Cre activity (Branda and Dymecki, 2004; Carney and Mosimann, 2018; Sauer and Henderson, 1988). Fusion of Cre with effector domains that enable chemical control of its activity, in particular with the ligandbinding domain of the Estrogen Receptor (ER) and its tamoxifen-responsive version T2 as Cre-ERT2 has resulted in further temporal control of recombinase activity (Feil et al., 1996; Metzger et al., 1995). Zebrafish are ideally suited for spatio-temporally controlled CreERT2 experiments, as the immediately active 4-OH-Tamoxifen (4-OHT) can simply be added to the embryo medium, resulting in detectable *loxP* reporter recombination within 1-2 h (Carney and Mosimann, 2018; Felker et al., 2016; Hans et al., 2009; Hans et al., 2011; Mosimann et al., 2011). Lineage tracing using *drl*-driven CreERT2 recombinase at early gastrulation stages enables Cre/*lox*-based lineage labeling of LPM-derived fates, including of virtually all myocardium (Felker et al., 2018; Mosimann et al., 2015; Prummel et al., 2019). The gradual refinement of *drl* reporter expression to FHF descendants in the heart provides an opportunity to preferentially label this progenitor pool and follow their contribution to the adult zebrafish heart.

Here, we document the correlative *drl*-based myocardial labeling of FHF descendants in the embryonic and in the adult zebrafish ventricle. Expanding previous findings, late gastrulation-stage labeling using *drl:creERT2* with a *myl7*-controlled, myocardium-selective *loxP* transgene results in predominant labeling of FHF-derived outer curvature (OC) and the right side of the embryonic ventricle. Raised to adulthood, such lineage-labeled hearts featured only rarely labeled cells in the atrium, yet the labeling in the ventricle is confined to a part of the ventricle that is positioned at the opposite side to the atrium. These seemingly FHF-descending ventricle cardiomyocytes form a clear boundary to the myocardium in the absence of any physical boundary. Our data documents a persisting cell-based compartmentalization of the adult zebrafish ventricle proceeding from early cardiac development.

## Materials and Methods

### Zebrafish husbandry and procedures

All animal husbandry and procedures were carried out as approved by the cantonal veterinary office of Fribourg, Switzerland; the cantonal veterinary office of Zurich, Switzerland; and veterinary office of the IACUC of the University of Colorado School of Medicine, USA (protocol #00979).

### Lineage labeling and analysis of embryonic hearts

To perform myocardial lineage labeling, heterozygous male *Tg(drl:creERT2; cryaa:Venus)^cz3333Tg^* (Mosimann et al., 2015) (abbreviated as *drl:creERT2* throughout the manuscript) were individually crossed to heterozygous female *Tg(myl7:LOXP-AmCyan-LOXP-ZsYellow)^fb2Tg^* (Zhou et al., 2011) (abbreviated as *myl7:Switch*). Collected embryos were kept in E3 medium at 28°C for the duration of the experiments. Cre/lox experiments were performed according to our previous reports and guidelines (Felker and Mosimann, 2016; Felker et al., 2016). In detail, CreERT2 activity was induced at 75% epiboly with E3 containing 10 μM final concentration of (Z)-4-Hydroxytamoxifen (Sigma Aldrich, H7904, abbreviated as 4-OHT). 4-OHT stock was dissolved in DMSO as 10 mM single-use aliquots stored at −20°C in the dark, and 4-OHT aliquots were used up within 1-2 months of dissolving as stock solution (Felker et al., 2016). 4-OHT treatment was performed overnight. After treatment, 4-OHT was washed out of the embryo medium and replaced with N-Phenylthiourea (Sigma Aldrich, P7629) at a final concentration of 75 μM in E3 embryo medium to inhibit melanogenesis for embryos to be imaged at 3 dpf. Animals pursued for later time points were raised in E3 and transferred to nursery tanks after 5 dpf.

At 3 dpf, embryos were anesthetized with 0.016% Tricaine-S (MS-222, Pentair Aquatic Ecosystems, NC0342409) in E3 embryo medium and embedded in E3 with 1% low-melting-point agarose (Sigma Aldrich, A9045) with 30 mM 2,3-butanedione monoxime (Sigma Aldrich, B0753) to stop heartbeat during imaging. Embryos were mounted on glass bottom culture dishes (Greiner Bio-One, 627861) orienting the anterior dorsal side of the embryo toward the bottom of the plate. Confocal imaging of the embryonic heart was performed with a Zeiss LSM880 using a × 20/0.8 air-objective lens. The blue (AmCyan) and yellow (ZsYellow) channels were acquired sequentially with maximum speed in bidirectional mode. The range of detection for each channel was adapted to avoid any crosstalk between the channels. Images of acquired Z-stacks were reconstructed with Fiji software (Schindelin et al., 2012) as a maximum intensity projection.

The percentage of ZsYellow-positive cells was quantified using the Fiji 3D Objects Counter plugin on each acquired Z-projection (Schindelin et al., 2012). The threshold was set to 30 voxels and maintained across each embryo. Equidistant lines were drawn across the center of the atrium and ventricle to define the midpoint and distinguish ‘left’ and ‘right’ of each structure (Fig. 1). The ROI of each structure (left atrium, right atrium, left ventricle, right ventricle, respectively) was defined and the region outside of the ROI was cleared. Channels were then split into blue and yellow and quantified using the 3D Objects Counter Plugin. Object voxels from each channel (blue or yellow, respectively) were compared to total object voxels from each ROI. Error bars correspond to standard error of the mean (SEM). Significance of differences was calculated using two-tailed Student’s t-test or one-way ANOVA with Tukey’s multiple comparison test. Statistical analyses were performed with GraphPad Prism. All results are expressed as the mean ± SEM.

**Figure 1.**
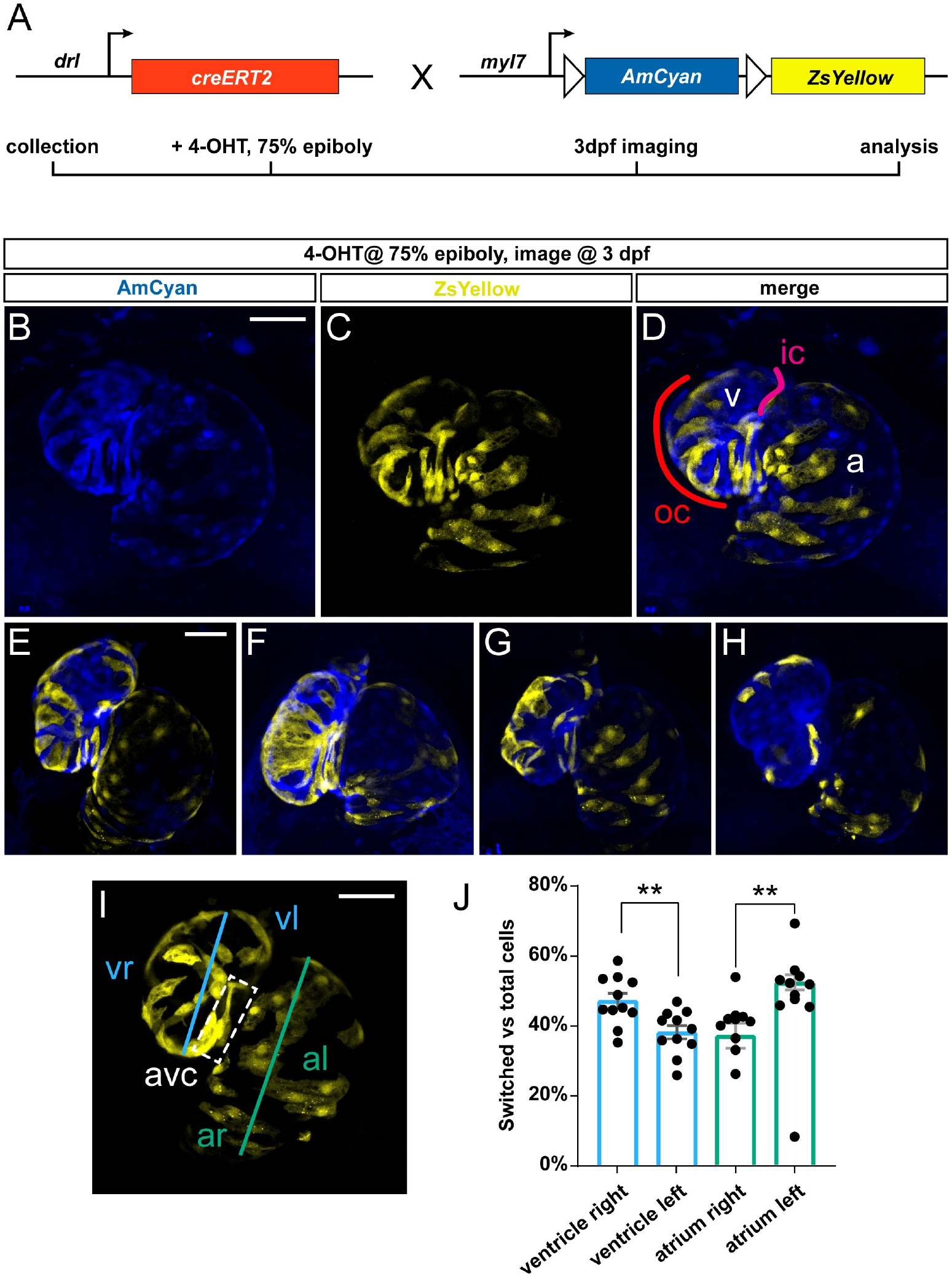
Predominant labeling of the FHF-assigned embryonic myocardium using *drl:creERT2*. (**A**) Crossing scheme of used transgenes. *drl:creERT2* is expressed in the gastrulation-stage LPM progenitors and becomes gradually restricted to FHF descendants in the heart, while *myl7:Switch* is a cardiomyocyte-specific *loxP* reporter line that by default marks myocardium in AmCyan (blue) and ZsYellow (yellow) upon exposure to active Cre recombinase. Timeframe of involved experimental steps outlined below. See text for details. (**B-H**) Maximum intensity projections of confocal stacks taken from 3 dpf zebrafish embryos double-transgenic for *dr:creERT2;myl7:Switch* and 4-OHT-induced at 75% epiboly. Ventral views, rostral to the top. Individual color channels and merge shown for an individual heart (**B-F**), atrium (a) and ventricle (v) annotated for predominantly SHF-derived inner curvature (ic), predominantly FHF-derived outer curvature (oc). Additional hearts of the imaging series shown below (**E-H**), note the distribution of *drl:creERT2*-induced ZsYellow clones throughout atrium and ventricle. (**I,J**) Quantification of clone distribution throughout atrium and ventricle, division lines for left vs right atrium (al, ar) and ventricle (vr, vl), with boxed atrioventricular canal shown (**I**) that predominantly lies on the left side using this analysis and contributes to possible overrepresentation of left-sided clones in the ventricle. p=0.002 for atrium, p= 0.0046 for ventricle, bar diagram depicts mean with error bars as SEM, significance based of differences was calculated using two-tailed Student’s t-test (n=11 hearts). Scale bars: 50 μm.

### Imaging and analysis of adult hearts and heart sections

Immunofluorescence analyses of heart sections were performed essentially as previously described (Bise et al., 2020). Briefly, entire larvae or dissected adult hearts were fixed in 2% paraformaldehyde overnight at 4 °C. After wash in PBS, specimens were equilibrated in 30% sucrose at 4 °C, embedded in tissue freezing media (Tissue-Tek O.C.T., Sakura) and cryosectioned at a thickness of 16 μm on Superfrost Plus slides (Fisher). Embryos and larvae were sectioned along the coronary body axis, whereas adult hearts were cut transversally from the bulbus arteriosus towards the apex. Slides were stored at −20°C. Before use, the slides were rehydrated in 0.3% Triton-X in PBS (PBST) and incubated in blocking solution (5% goat serum in PBST) for 1 h at RT. Primary antibody mouse anti-Tropomyosin (developed by J. Jung-Chin Lin and obtained from Developmental Studies Hybridoma Bank, CH1) was diluted in blocking solution at 1:100 and was applied on the sections overnight at 4°C. Slides were washed in PBST and incubated in secondary antibodies (1:500, Jackson ImmunoResearch Laboratories) in blocking solution for 2 h at RT. After washing in PBST, slides were mounted in 90% glycerol in 20 mM Tris pH 8 with 0.5% N-propyl gallate.

Fluorescent images of sections were taken with a Leica TCS SP5 confocal microscope, and ImageJ 1.49c software (Schneider et al., 2012) was used for subsequent measurements. For quantification of *drl:creERT2;myl7:Switch*-labeled cardiomyocytes after 4-OHT induction, we calculated the proportion of ZsYellow-positive area superimposed with the Tropomyosin-positive area per ventricle. For this, the ROI (atrium, atrium-proximal ventricle and atrium-distal ventricle) was selected. Channels were split and then blue and yellow channels were superimposed using Colocalization Plugin (ratio 40%, threshold 50.0). The area of positive signal was measured using threshold and compared with the area of Tropomyosin (blue channel). Error bars correspond to standard error of the mean (SEM). Significance of differences was calculated using two-tailed Student’s t-test or One-Way ANOVA with Tukey’s multiple comparison test. Statistical analyses were performed with GraphPad Prism. All results are expressed as the mean ± SEM.

## Results

### Preferentially labeling of first heart field-derived myocardium using drl:creERT2

We previously established that *drl:creERT2* combined with *loxP* reporters broadly labels LPM and all cardiac lineages when induced in early gastrulation (shield stage, 6 hpf), while 4-OHT-triggered labeling increasingly specifies in the myocardium to the FHF descendants that at 72 hpf encompass the OC on the predominantly right side of the embryonic ventricle (Felker et al., 2018; Mosimann et al., 2015; Sánchez-Iranzo et al., 2018). Towards establishing how the FHF descendants contribute to the myocardium over time, we crossed *drl:creERT2* with the *myl7:loxP-AmCyan-STOP-loxP-ZsYellow* (referred to as *myl7:Switch*), which by default marks *myl7*-expressing cardiomyocytes with blue fluorescence and with yellow fluorescence after Cre/*lox-* mediated recombination with medium switching efficiency compared to other *loxP* reporters (Fig. 1A) (Felker et al., 2018; Mosimann et al., 2015; Zhou et al., 2011). To label *drl*-positive cells, we treated double-transgenic *drl:creERT;myl7:Switch* embryos at 75% epiboly with 4-OHT or DMSO solvent as control and imaged the resulting fluorescence pattern in the hearts at 3 dpf. In agreement with previous descriptions (Felker et al., 2018; Mosimann et al., 2015), we observed ZsYellow-fluorescent lineage labeling in both the ventricle and the atrium across their different regions (n=11) (Fig. 1B-H). Nonetheless, we noted that ventricular lineage labeling preferably occurred at the atrio-ventricular canal (AVC) and on the right side of the ventricle or OC, in line with the predominant FHF-derived origin of this myocardium domain (de Pater et al., 2009; Hami et al., 2011; Lazic and Scott, 2011; Mosimann et al., 2015).

Towards attaining an approximate quantitative measure of how *drl* lineage-labeled cells distribute in the ventricle, we analyzed recombined hearts for the ratio of switched (ZsYellow-fluorescent) to unswitched (AmCyan-fluorescent) cells in the right versus the left side of the ventricle (Fig. 1I,J) (see Methods for details). Of note, due to the twisted morphology of the ventricle and atrium, our left versus right measurement is a mere approximation, likely underestimating the contribution to the right side (Fig. 1I). Nonetheless, our analysis revealed that, though variable, a significant portion of recombined ventricle cells resided in the OC and right side versus the inner curvature (IC) on the left side (Fig. 1J) (n=11). While *drl:creERT2* faithfully labels myocardium contributing to all parts of the ventricle in accordance with its LPM origin, our analysis is in line with, and expands upon, the notion that the activity of *drl*-based transgenic reporters progressively refines to the FHF progenitors (Mosimann et al., 2015).

To extend our analysis to later developmental stages, we performed coronal sections of *drl:creERT;myl7:Switch* larvae at 5 dpf and 22 dpf that were induced at 75% epiboly with DMSO solvent or 4-OHT (Fig. 2A). To demarcate the cardiac tissue on sections, we used Tropomyosin immunostaining, which also detects skeletal muscles (Pfefferli and Jaźwińska, 2017). The hearts of DMSO-treated control zebrafish displayed AmCyan expression, but not ZsYellow, consistent with tightly controlled CreERT2 activity (Fig. 2B). In 4-OHT-treated zebrafish, we observed at 5 and 22 dpf that the ventricular myocardium retained ZsYellow lineage label (Fig. 2C-D) (5 dpf n=6; 22 dpf, n=6). Quantification of the ZsYellow-labeled area within the cardiac tissue, as demarcated by Tropomoysin immunostaining (marking all cardiomyocytes), documented approximately 28-52% switched ventricular cardiomyocytes at both timepoints (Fig. 2E), indicating that the lineage-labeled myocardium continues to substantially contribute to the growing ventricle. Taken together, *drl:creERT2-based* lineage labeling using *myl7:Switch* preferentially labels myocardium assigned to a FHF origin and this labeling is retained during later developmental stages.

**Figure 2.**
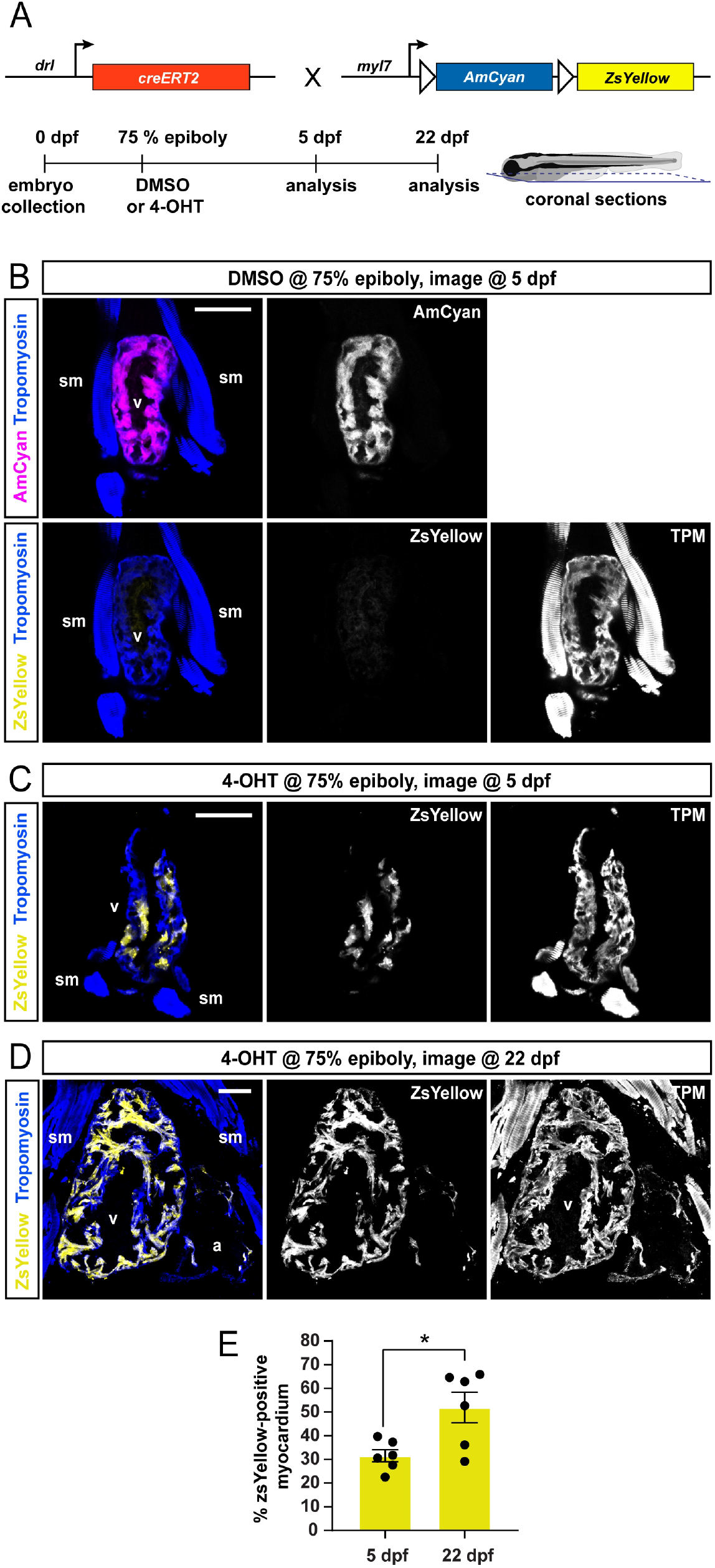
FHF lineage labeling persists during developmental stages. (**A**) Crossing scheme of used transgenes and timeline of experimental setup with 5 dpf embryos and 22 dpf larvae longitudinally sectioned for heart analysis. (**B-D**) Confocal images of individual coronal sections, stained for pan-muscle Tropomyosin (blue, sm indicating skeletal muscles, v indicating ventricle, a indicating atrium). Embryos at 5 dpf (**B**), treated with DMSO (control) at the 75% epiboly, show only myocardial AmCyan expression from *myl7:Switch* (n=4), while 4-OHT treatment (**C**) results in ZsYellow-expressing clones from *drl:creERT2* activity, as also observed at 22 dpf (**D**). (**E**) Quantification of the ZsYellow-Tropomyosin double-positive cardiomyocytes of analyzed ventricles, two-tailed Student’s t-test, * p=0,0149, 5 dpf n=6; 22 dpf n=6. Scale bar 50 μm.

### FHF cardiomyocytes enrich at the atrium-opposite side of the adult ventricle

To examine the distribution of *drl*-based FHF lineage labeling in the adult zebrafish myocardium, we again induced *drl:creERT2;myl:Switch* fish with 4-OHT at 75% epiboly stage and raised them to adulthood (Fig. 3A). At the age of 4 months, we performed transversal sections of fixed hearts starting from the ventricle base towards the apex (Fig. 3B). We then analyzed ZsYellow expression from recombined *myl7:Switch* in the Tropomyosin-stained myocardium in serial sections. Consistent with our developmental observations (Fig. 1, 2), the adult myocardium remained partially labeled by ZsYellow expression (n=8, two experimental replicates) (Fig. 3C-H). Notably, also in our adult heart sections, the distribution of ZsYellow-fluorescent cardiomyocytes was not randomly spread through the myocardium, but displayed an asymmetric pattern across cardiac chambers. Throughout, the analyzed atria only contained few ZsYellow-labeled cells, while the ventricles consistently harbored broader areas of ZsYellow-labeled cells (Fig. 3C-H). These data indicate that the FHF-dominating lineage labeling we observed at early developmental stages translates to regionalized cardiomyocyte labeling in the adult zebrafish hearts.

**Figure 3:**
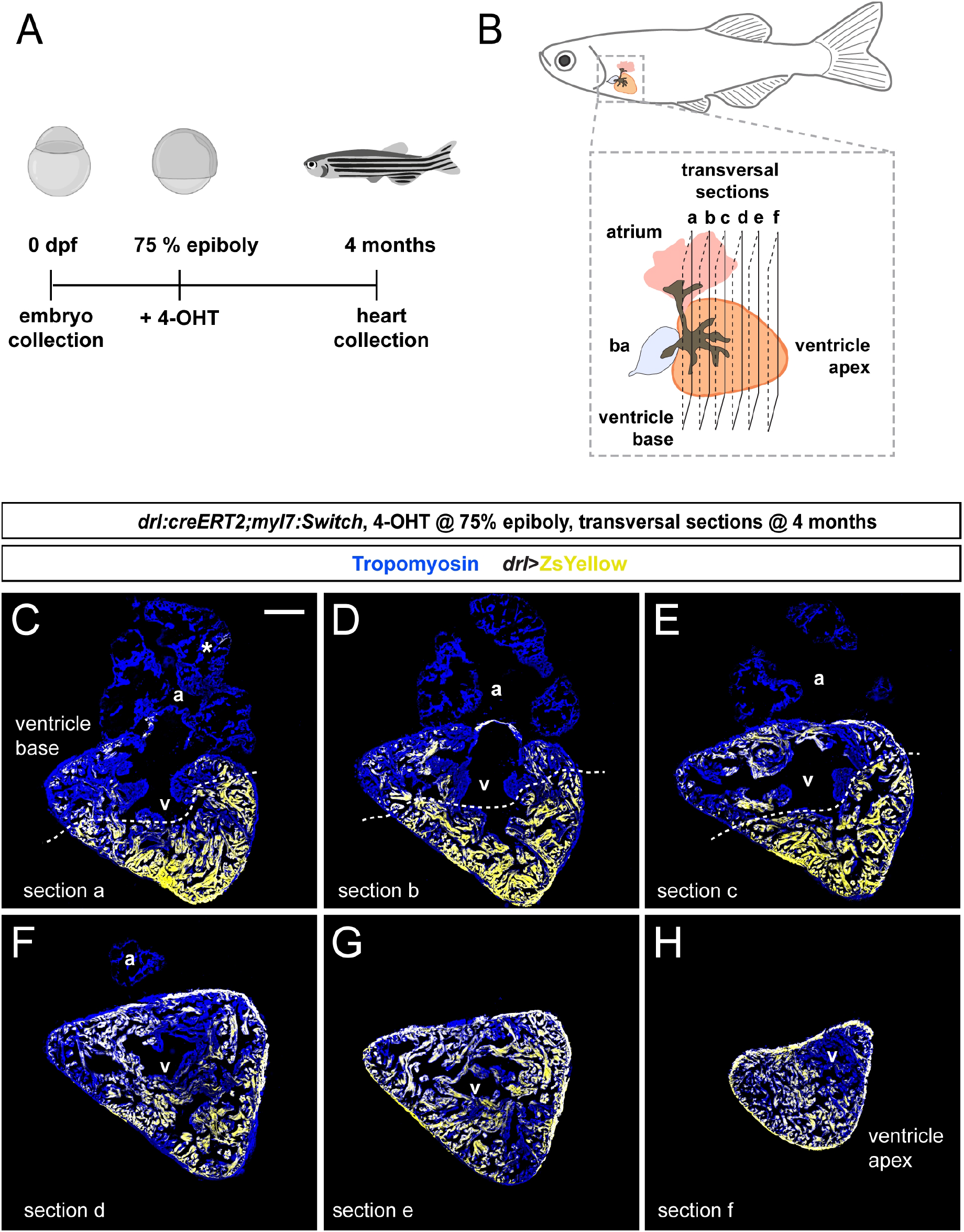
FHF contribution to the adult zebrafish ventricle remains localized. (**A-B**) Schematic of the experimental design and timeline. Embryos were 4-OHT-treated at 75% epiboly, raised to adulthood for 4 months. Hearts were dissected from the adult animals (**B**) sequentially sectioned from the base of the heart to the apex as per schematic. (**C-H**) Series of transversal sections of a selected heart, immunostained against Tropomyosin (blue) and showing ZsYellow expression (yellow) from *drl:creERT2*-recombined *myl7:Switch* with atrium (a) on top and ventricle (v) at the bottom. Asterisk denotes rare atrial ZsYellow clones throughout sections. Note the markedly larger area of ZsYellow-labeled cardiomyocytes in the ventricle. Scale bar: 200 μm.

To gain a quantitative and regionalized measurement of the FHF-derived cardiomyocyte contribution to the ventricle, we selected the two largest transversal sections at the ventricle base and just below this region of each heart (n=8) (Fig. 4A). The sampled sections represented non-adjacent regions separated by approximately 100 μm. From this analysis, we found that the sectioned atria contained on average 4% of ZsYellow-positive cardiomyocytes (Fig. 4B-D). Following the asymmetrical ZsYellow labeling on ventricle sections, we accordingly subdivided the tissue into two zones of high versus low label retention, respectively. Although our approach entails approximation, we determined that the myocardium with low-level ZsYellow labeling (27%) predominantly located proximal to the atrium and includes the ventricle base, whereas over double the amount of Zs-Yellow labeling occurred in the region distal to the atrium that also includes the apex (69% of ZsYellow-positive myocardium) (Fig. 4C-D). The overall pattern thus emphasizes that the atrium-distant portion of the ventricle chamber harbors predominantly FHF-descendant cardiomyocytes. Taken together, our data are consistent with a persistent separation of FHF and SHF myocardium that starts during embryogenesis and continues throughout the life of the zebrafish heart, resulting in a lineage-based compartmentalization of the adult zebrafish ventricle even in the absence of any septation (Fig. 5).

**Figure 4:**
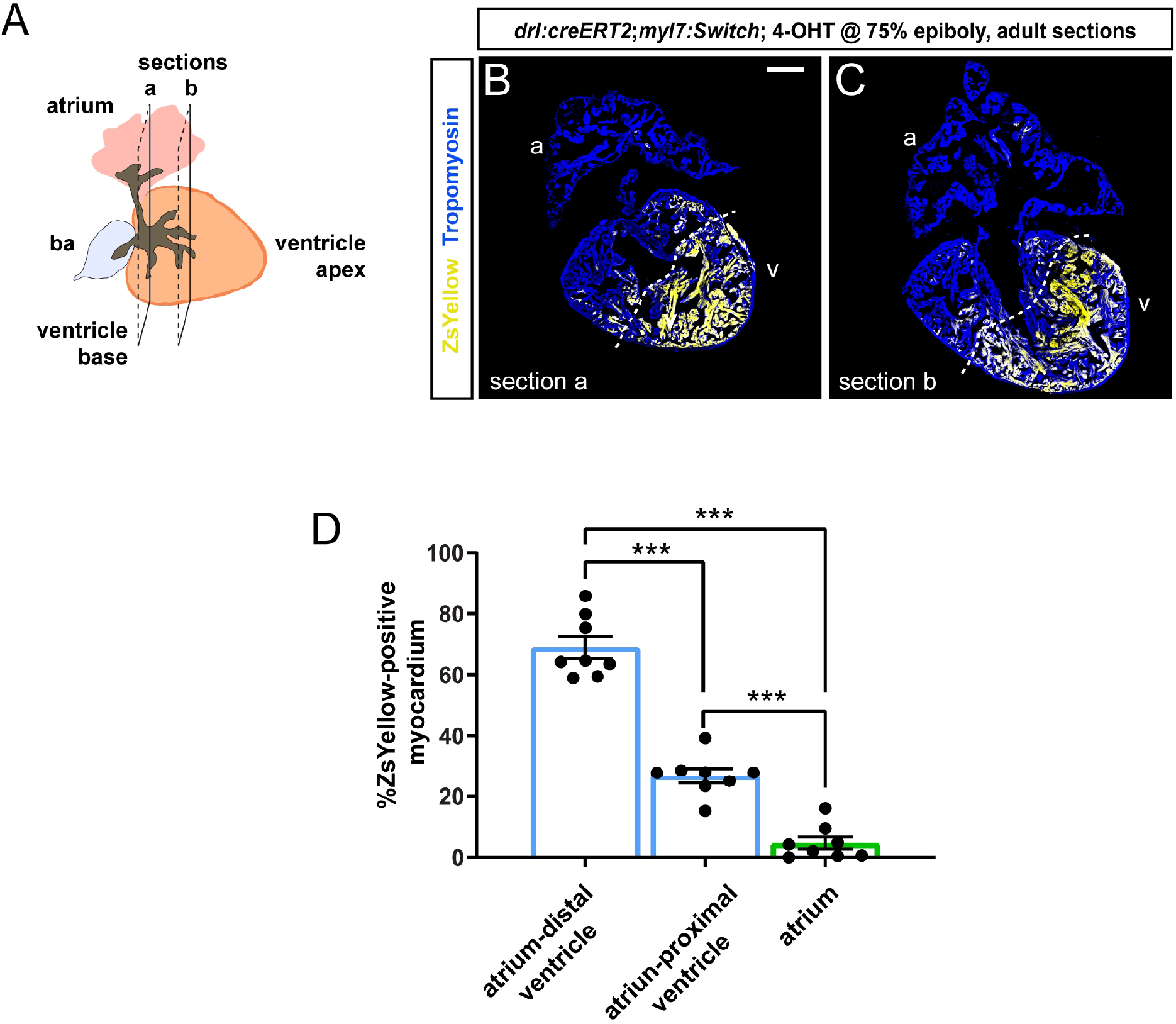
Quantification of localized FHF lineage labeling in the adult zebrafish ventricle. (**A**) Schematic illustration of the adult zebrafish heart with two planes of transversal sections across the ventricle and the atrium as analyzed. (**B, C**) The two heart sections (a, b) with largest tissue area were selected for quantification of ZsYellow-labeled cardiomyocytes. Dotted line depicts possible boundary of high-versus low-density ZsYellow labeling. Note that nonetheless no sharp boundary separates the two areas. Scale bar: 200 μm. (**D**) Quantification of ZsYellow-positive cardiomyocyte area in the Tropomyosin-demarcated myocardium indicates predominant labeling in the apex-distal ventricle that includes the apex, while the region of the ventricle base only retained low numbers of ZsYellow-labeled cells. Note the scarce atrium labeling. One-Way Anova test followed by Tukey’s multiple comparison test, *** p<0.0001 (n=8 hearts, 2 sections each as per **A**).

**Figure 5.**
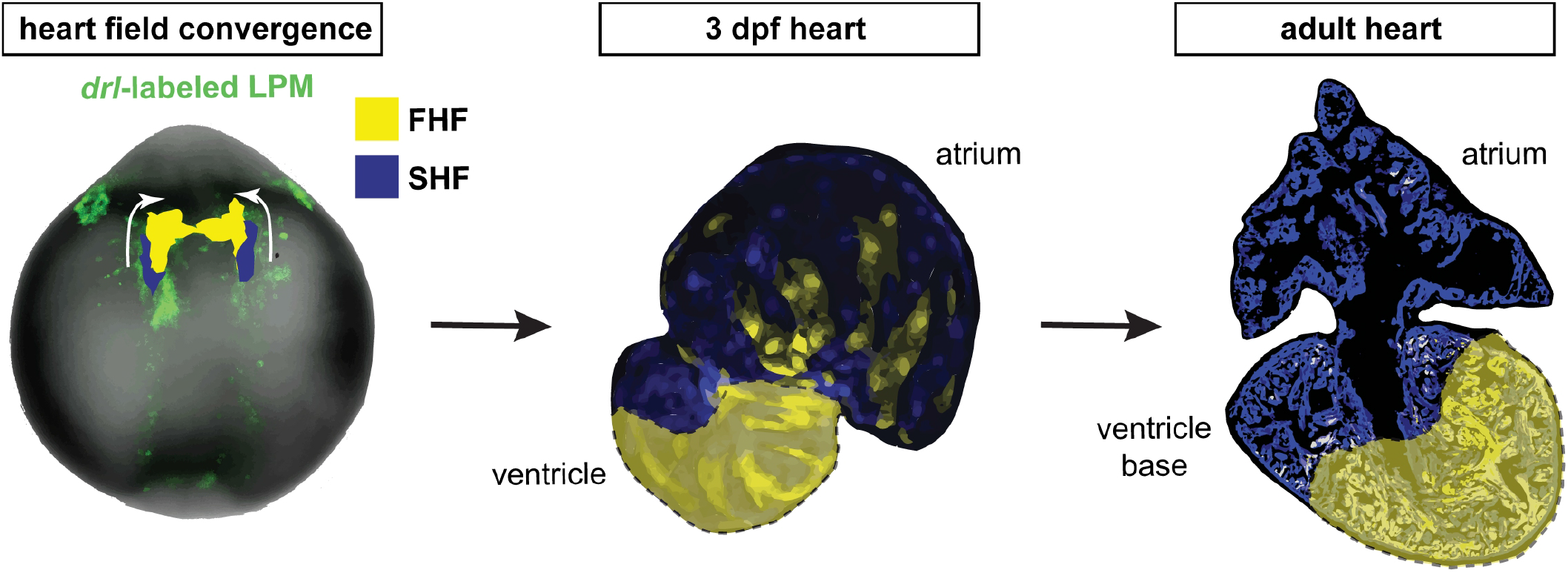
*drl-based* FHF lineage tracing of cardiomyocytes reveals compartmentalization of the adult zebrafish ventricle. Model summary of the reported findings. FHF and SHF descendants in the ALPM form the early-differentiating and late-differentiating myocardium, respectively, in the early heart. In the adult heart, the FHF myocardium then predominantly contributes to the atrium-distal portion of the ventricle including the apex, whearas the atrium-proximal region of the ventricle base is predominantly SHF-derived. The interface between the FHF and SHF regions is nonetheless not a sharp compartment boundary.

## Discussion

Developmental integration of FHF and SHF lineages is a seemingly universal trait among vertebrate hearts. The mammalian ventricle splits into a left and a right chamber by forming a septum between the FHF and the SHF descendants of the ventricle myocardium. This physical division is central to the formation of a separate systemic and pulmonary circuit. Evolutionarily adapted to and representative of teleost physiology, the zebrafish ventricle directly moves blood to the gill arches as part of a single systemic circuit, yet also integrates FHF and SHF myocardium in incompletely understood dynamics and pattern. Here, we present data that support and extend previous work that established cell-level compartmentalization of the zebrafish ventricle in the absence of physical septation.

Using *drl* transgenic-based lineage tracing that marks all myocardium, but predominantly FHF-descending cardiomyocytes in our used framework (Mosimann et al., 2015), we document the persistent distinction of FHF versus SHF contribution to the zebrafish ventricle from the embryonic to the adult heart. In the latter, the FHF predominantly showed contribution to the atrium-distant regions of the ventricle including the apex (Fig. 3, 4). This finding is in line with our previously reported lineage tracing using *tbx5a*-based lineage tracing that seemingly encompasses and expands beyond the *drl*-based labeling (Sánchez-Iranzo et al., 2018). Indeed, *drl:creERT2* with our 4-OHT induction regimen seems to label a more restricted region of the ventricle (Fig. 4, 5). These partial discrepancies between *tbx5a*- and *drl*-based lineage analyses could stem from several factors. First, the used cardiomyocyte-specific *loxP* reporter *myl7:Switch* might be more selectively permissive for recombination than other *loxP*-based reporter lines used previously with *drl:creERT2* including *ubi:Switch* and *hsp70l:Switch*, such as for high levels of CreERT2 that *drl*-based reporters build up over time in FHF progenitors (Felker et al., 2018; Mosimann et al., 2015; Prummel et al., 2019; Sánchez-Iranzo et al., 2018). Second, the histological orientation of hearts presented in different studies, specifically selecting a sagittal versus transversal sectioning plane, reveals different anatomical perspectives of the organ that emphasizes distinct spatial distribution of labeled cells. Nonetheless, while *tbx5a*-based lineage labeling to the adult ventricle appears to mark a broader territory than *drl*-based labeling (Sánchez-Iranzo et al., 2018), both lineage tracing analyses emphasize 1) the region of the ventricle base forms from FHF and SHF myocardium, while 2) the region centered around the apex is predominantly of FHF descent. Further comparison of *tbx5a* versus *drl* and other myocardium-labeling CreERT2 drivers is warranted to elucidate that the *bona fide* FHF territory in the adult ventricle represents.

Previous work in mouse has elegantly documented how the SHF descendants form the right ventricle and contribute to the ventricle-separating septum, such as tracking by the *Mef2c* AHF enhancer-driven Cre (Dodou et al., 2004; Verzi et al., 2005). Establishment of a gradient of Tbx5 expression in tetrapod evolution has been causally linked to septum formation between these FHF and SHF descendants in the ventricle (Koshiba-Takeuchi et al., 2009; Mori and Bruneau, 2004). While forming potent trabeculation, the single ventricle in teleosts is devoid of any septation. Curiously, in addition to ventricle asymmetry at a cellular and gene expression level, also the zebrafish atrium shows such apparent regionalization, as revealed by asymmetric *meis2b* expression in the left side of the adult atrium (Guerra et al., 2018). Our work presented here further supports this concept and highlights that the cellular subdivision of the ventricle persists along its distance from the atrium (Fig. 5). Nonetheless, we note that the distinction between FHF and SHF myocardium in the adult ventricle does not form a sharp compartment boundary and future studies are warranted to elucidate the molecular interactions that underlie the persistent lineage separation.

With increasing evidence for an asymmetrical contribution of diverse developmental lineage origins to the adult zebrafish heart, the functional contribution of this fascinating phenomenon to adult heart patterning and electrophysiology requires further investigation. Open questions remain whether other heart-associated systems including cardiac innervation and the coronary vasculature interact with, or provide any hallmarks associated with, a cardiac chamber subcompartmentalization during heart morphogenesis. The formation of FHF and SHF lineages is of ancient origin within the LPM, dating back to our early chordate ancestors (Diogo et al., 2015; Prummel et al., 2020; Stolfi et al., 2010). The functional aspect of persistent FHF versus SHF compartmentalization in the adult zebrafish heart hints at the existence of such continued separation already in the last common ancestor of teleosts and tetrapods, providing a versatile scaffold to adapt the circulatory system to terrestrial versus aquatic habitats over millions of years.

## Author Contributions

Conceptualization, C.M. and A.J.; Formal analysis, C.P., H.R.M. and A.F.; Visualization, C.P. and H.R.M.; Resources, C.M. and A.J.; Writing—original draft preparation, C.M. and A.J.; Writing—review and editing, C.P. and H.R.M.; Supervision, C.M. and A.J.; Project administration, C.M. and A.J.;Funding acquisition, C.M. and A.J. All authors have read and agreed to the published version of the manuscript.

## Institutional Review Board Statement

The study was conducted according to the guidelines of the Declaration of Helsinki and approved by the cantonal veterinary office of Fribourg, Switzerland, the cantonal veterinary office of Zurich, Switzerland, and the veterinary office of the IACUC of the University of Colorado School of Medicine (protocol #00979), USA.

## Acknowledgments

We thank Verena Zimmermann, and Christine Archer and Molly Waters for zebrafish facility management and husbandry, Dr. Caleb Doll for confocal imaging support, and the members of the Mosimann and Jaźwińska labs for input on the manuscript.

